# A *Histoplasma capsulatum* lipid metabolic map identifies antifungal targets

**DOI:** 10.1101/2020.03.02.973412

**Authors:** Daniel Zamith-Miranda, Heino M. Heyman, Meagan C. Burnet, Sneha P. Couvillion, Xueyun Zheng, Nathalie Munoz, William C. Nelson, Jennifer E. Kyle, Erika M. Zink, Karl K. Weitz, Kent J. Bloodsworth, Geremy Clair, Jeremy D. Zucker, Jeremy R. Teuton, Samuel H. Payne, Young-Mo Kim, Morayma Reyes Gil, Erin S. Baker, Erin L. Bredeweg, Joshua D. Nosanchuk, Ernesto S. Nakayasu

**Author notes:** These authors contributed equally (Daniel Zamith-Miranda, Heino M. Heyman, Meagan C. Burnet).

## Abstract

Lipids play a fundamental role in fungal cell biology, being essential cell membrane components and major targets of antifungal drugs. A deeper knowledge of lipid metabolism is key for developing new drugs and a better understanding of fungal pathogenesis. Here we built a comprehensive map of the *Histoplasma capsulatum* lipid metabolic pathway by incorporating proteomic and lipidomic analyses. We performed genetic complementation and overexpression of *H. capsulatum* genes in *Saccharomyces cerevisiae* to validate reactions identified in the map and to determine enzymes responsible for catalyzing orphan reactions. The map led to the identification of both the fatty acid desaturation and the sphingolipid biosynthesis pathways as targets for drug development. We found that the sphingolipid biosynthesis inhibitor myriocin, the fatty acid desaturase inhibitor thiocarlide and the fatty acid analog 10-thiastearic acid inhibit *H. capsulatum* growth in nanomolar to low micromolar concentrations. These compounds also reduced the intracellular infection in an alveolar macrophage cell line. Overall, this lipid metabolic map revealed pathways that can be targeted for drug development.

## INTRODUCTION

Fungal diseases affect more than 1 billion people and cause 1.6 million deaths every year (1). *Histoplasma capsulatum*, the causative agent of histoplasmosis, is an important pathogen in this context, being associated with HIV infections and causing morbidity and mortality worldwide (2). Data from 2011-2014 identified 3,409 cases of histoplasmosis in 12 states in the US with a 7% mortality rate (3). Serological surveys have shown reactivity against the *H. capsulatum* antigen histoplasmin in 60-90% of individuals in communities surrounding the Mississippi and Ohio basins, suggesting that the epidemiological data are underestimated (4,5). Histoplasmosis treatment relies on only a few antifungals and the increasing number of resistant strains is a major public health concern (6,7). The absence of immunotherapies and the fact that the newest class of antifungal drugs, echinocandins, being ineffective against *H. capsulatum*, make the development of new therapies a high priority. A major hurdle in developing new drugs is the limited knowledge about the detailed metabolic reactions of this microbe.

Lipids have essential roles in many biological processes and the biosynthetic pathways of fungal lipids diverged from metazoan pathways, which makes them obvious antifungal drug targets (8). Indeed, two classes of current antifungal drugs target lipid metabolism: 1) polyenes, including amphotericin B and nystatin, bind and extract ergosterol from the fungal membrane, respectively (9-11); and 2) azoles, such as voriconazole, itraconazole and fluconazole, inhibit cytochrome P-450-dependent 14α-sterol demethylase, an enzyme of the ergosterol biosynthetic pathway (8,12). More recently, glucosylceramide, a type of sphingolipid that is critical for infection of many fungal species, has been validated as a drug target in *Cryptococcus neoformans*, *Aspergillus fumigatus*, *Candida auris* and *Sporothrix* spp. (13-17).

There is limited information about *H. capsulatum* lipid composition and function, which is also true for most pathogenic fungi. Polyenes and azoles have been used to treat most mycoses, but many of these fungal species might not even produce ergosterol. Some species produce cholesterol, brassicasterol, lanosterol and other sterols as final products (18-22). However, it is still unclear how the different sterols affect the efficacy of polyenes and azoles. Considering the importance of lipids, we reasoned that a deeper characterization of *H. capsulatum* lipid metabolism would result in the discovery of drug targets. In this study, we performed an in-depth characterization of the *H. capsulatum* lipid biosynthetic pathway by profiling its lipids and the associated proteins, which were incorporated into a metabolic map. By comparing our results to *Saccharomyces cerevisiae* and humans, we show unique features of the lipid biosynthetic pathway of *H. capsulatum* that can be targeted for drug development.

## RESULTS

### Proteomic and lipidomic analyses and overview of the *H. capsulatum* lipid metabolism

To determine the global landscape of the lipid metabolic pathway, we performed comprehensive lipidomic and proteomic analyses of log-phase yeast-form *H. capsulatum* grown in Ham’s F12 medium. We chose this medium since it is a defined medium, allowing us to distinguish lipids that are synthesized by the fungus vs. the ones obtained from nutrients. Total lipids were extracted using two different methods and submitted to chromatographic separation before analyses by mass spectrometry. Paired global lipidomics and proteomics analyses were performed by submitting the samples to a simultaneous metabolite, protein and lipid extraction (MPLEx) (23) (**FIGURE 1A**). For complementary analyses of sterols, free fatty acids and phospholipids, yeast cells were extracted with two rounds of organic solvent extraction, followed by a solid phase fractionation in a silica 60 column (**Figure 1A**). The extracted fractions were analyzed by either gas chromatography-mass spectrometry (GC-MS) (sterols and fatty acids) or liquid chromatography-tandem mass spectrometry (LC-MS/MS) (sphingolipids, glycerolipids, phospholipids and proteins) (**Figure 1A**).

**Figure 1.**
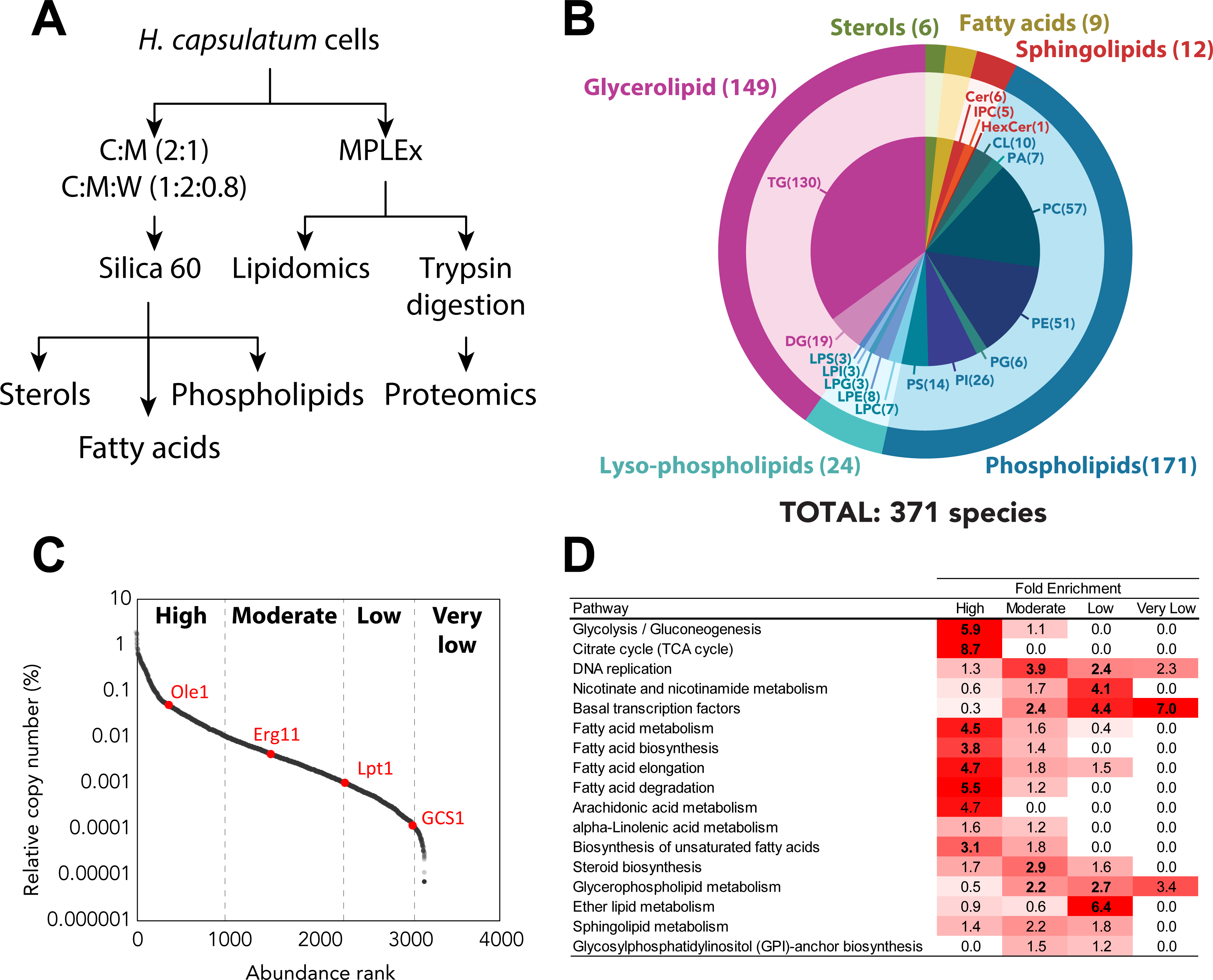
Proteomic and lipidomic analyses of *Histoplasma capsulatum* yeast cells. (A) Extraction procedure for lipidomics proteomics analyses. Yeast cells were extracted sequentially with chloroform:methanol (2:1, v:v) and chloroform:methanol:water (1:2:0.8, v:v:) and fractionated by Silica 60 solid phase extraction for sterol, free fatty acid and phospholipid analyses. Cells were also submitted to simultaneous metabolite, protein and lipid extraction (MPLEx) for global lipidomics and proteomics analyses. (B) Overall lipid coverage combining MPLEx and specific lipid extractions of *Histoplasma capsulatum*. (C) Protein abundance classification based on their relative copy numbers. (D) Function enrichment analysis based on the KEGG annotation of *H. capsulatum* proteins with different abundance levels. Abbreviations: C, chloroform; Cer, ceramide; CL, cardiolipin; DG, diacylglycerol; FA, fatty acid; HexCer, hexosylceramide; LPC, lysophosphatidylcholine, LPE, lysophosphatidylethanolamines, LPG, lysophosphatidylglycerol; LPI, lysophosphatidylinositol; LPS, lysophosphatidylserine; M, methanol; PA, phosphatidic acid; PC, phosphatidylinositol; PE, phosphatidylethanolamine; PG, phosphatidylglycerol; PI, phosphatidylinositol; PI_Cer, inositolphosphoceramide; PL, phospholipid; PS, phosphatidylserine; ST, sterol; TG, triacylglycerol; W, water.

The combined analysis identified 371 unique lipid species from 5 major lipid categories (fatty acids, sterols, glycerolipids, sphingolipids and glycerophospholipids (phospholipids + lysophospholipids)) that were subdivided into 19 subclasses (**Figure 1B** and **Tables S1-S8**). The most diverse subclasses of lipids in terms of the number of identified species were triacylglycerols (TG), phosphatidylcholines (PC) and phosphatidylethanolamine (PE) with 130, 57 and 51 individual species, respectively (**Figure 1B**). The proteomic analysis led to the identification of 3,215 proteins (**Table S9**).

To provide a measurement of the protein abundances in the cells, we calculated the relative copy number of each protein and scaled them into high, moderate, low and very low abundance (**Figure 1C**), using a scale similar to one previously described (24). To validate this scale, we performed a function-enrichment analysis using the KEGG annotation to check the abundance of different pathways. As expected, glycolysis/gluconeogenesis and tricarboxylic acid (TCA) cycle were overrepresented among the highly abundant proteins, whereas DNA replication, nicotinate and nicotinamide metabolism, and basal transcription factors were enriched in moderate, low and very low abundance levels, respectively (**Figure 1D**). The same scale showed that fatty acid metabolism was enriched in highly abundant proteins, whereas steroid biosynthesis proteins were mainly present in moderate abundance (**Figure 1C-D**). Glycerophospholipid and sphingolipid metabolism proteins were not concentrated in a single abundance level and were spread mostly between moderate and low abundant levels (**Figure 1C-D**). The abundance levels of the proteins were directly proportional to each type of lipid in the cellular membrane. For instance, the proteins of fatty acid and steroid (the building blocks of the cell membrane; therefore, most abundant lipids) metabolism were significantly enriched among the high and moderate abundance proteins (highlighted with bold font in the first two columns of **Figure 1D**). Glycerophospholipid, ether lipid or sphingolipid metabolism, which are responsible for the synthesis of lipids from specific lower-abundance classes, were enriched in proteins with moderate to very low abundance proteins (**Figure 1D**).

### Lipid biosynthesis and remodeling map of *H. capsulatum*

To provide a global view of *H. capsulatum* lipid metabolism we built a map including lipid biosynthesis and remodeling reactions. The map was constructed based on a metabolic model (25) structured according to conserved metabolism between *H. capsulatum* vs. *S. cerevisiae*. We added information available for *Cryptococcus neoformans*, one of the best-characterized fungal organisms in terms of lipid metabolism (**Table S10**), along with previous literature on *H. capsulatum*. We used the *H. capsulatum* genomic, proteomic and lipidomic information to further restrict or add reactions that are present in this organism. The *H. capsulatum* map was incorporated with the relative abundance of the lipid species within the same subclass and the protein abundances (**Figure 2**). In terms of fatty acids, species containing 16 and 18 carbons were most abundant. Consistent with Figure 1D, 5 out of 10 proteins of this pathway were highly abundant (**Figure 2**). Similarly, 13 of the 23 sterol biosynthesis proteins had moderate abundance, with ergosterol being the most abundant product (**Figure 2**). In the sphingolipid pathway, ceramides (Cer), hexosylceramide (HexCer), and inositolphosphoceramides (PI_Cer) were the detected lipid species (**Figure 2**). Out of the 9 proteins detected in the proteome, 3 had low abundance, 5 had moderate abundance and 1 was highly abundant (**Figure 2**). The number of low abundant proteins does not necessary mean that the pathway has low activity. For instance, one low abundance protein had more abundant paralogues with the same function (serine palmitoyltransferases Lcb2 vs. Lcb1), and the other two proteins regulate the specific modification of the head group of HexCer, glucosylceramide synthase Gcs1 and endoglycoceramidase-related protein EGCrP1 (**Figure 2**). In terms of glycerolipids, diacylglycerols (DGs) and TGs were identified, being the proteins from this pathway that were present at moderate (2), low (3) and very low abundance (1) (**Figure 2**). Like the free-fatty acid composition, the most abundant species of DGs and TGs had fatty acyl groups with either 16 or 18 carbons attached to them (**Figure 2**). Consistent with the glycerolipids, all the different glycerophospholipid classes had species bearing C16 and C18 as the most abundant in each (**Figure 2**). To increase the utility of the lipid metabolic map, we developed the map into an informatics tool to visualize lipidomics and proteomics data. We tested the tool using lipidomics and proteomics data of *Candida albicans* from a previous publication (26), which showed a similar pattern of lipid species with fatty acids containing 16 or 18 carbons in length being the most abundant ones (**Figure S1**).

**Figure 2.**
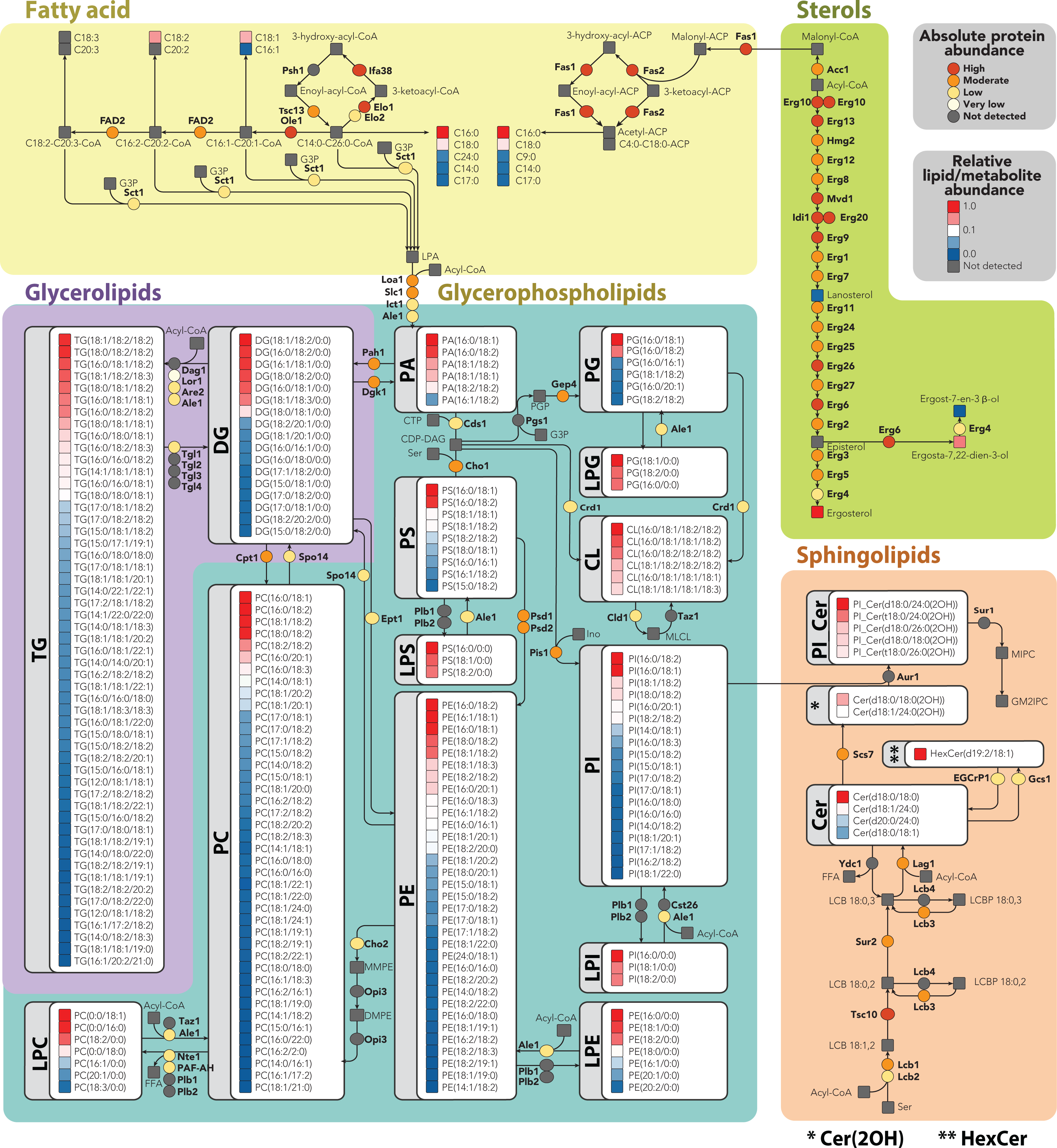
Metabolic map of *Histoplasma capsulatum* lipids and biosynthesis proteins. The map was built based on genomic information along with proteomics and lipidomics data. Lipids abundances were normalized by the most intense signal in the mass spectrometry analysis within each lipid class, whereas the proteins were quantified based on their relative copy number. For details about enzyme names and homologs see table S10. Abbreviations: Cer, ceramide; CL, cardiolipin; DG, diacylglycerol; HexCer, hexosylceramide; LPC, lysophosphatidylcholine, LPE, lysophosphatidylethanolamines, LPG, lysophosphatidylglycerol; LPI, lysophosphatidylinositol; LPS, lysophosphatidylserine; M, methanol; PA, phosphatidic acid; PC, phosphatidylinositol; PE, phosphatidylethanolamine; PG, phosphatidylglycerol; PI, phosphatidylinositol; PI_Cer, inositolphosphoceramide; PL, phospholipid; PS, phosphatidylserine; TG, triacylglycerol.

We further curated the map because homologous genes can have different specificities. We chose to determine the substrates and products of the *H. capsulatum* acyltransferase Ale1 (also known as lysophosphatidylcholine (LPC) acyltransferase 1 – LPT1) because its homolog in *S. cerevisiae* has been shown to modify different classes of lipids (27). This would allow coverage of a large portion of *H. capsulatum* lipid metabolism. Unfortunately, gene knockout is very difficult in *H. capsulatum* and RNAi experiments often result in a reduction of expression levels that fail to produce a phenotype. Conversely, exogenous expression of genes from the lipid metabolism in biotechnological fungi, such as *S. cerevisiae* and *Yarrowia lipolytica*, has provided important insights on enzyme specificity (28,29). Therefore, we performed complementation studies in *S. cerevisiae* with the *H. capsulatum Lpt1* (*Hc-Lpt1*), which would allow curation of a large portion of the metabolic map. A lipidomic analysis was performed in the *Lpt1* knockout strain of *S. cerevisiae* complemented or not with the *Hc-Lpt1* homolog (53% similarity to *S. cerevisiae Lpt1* – *Sc-Lpt1*) (**Figure S2**), using a plasmid with a galactose-inducible promoter. This allows specific expression of *Hc-Lpt1* in the presence of galactose, but not glucose. Both gene knockout and recombinant expression were validated by proteomic analysis (**Figure 3A**). As expected, the Sc-LPT1 specific peptide DISASSPNLGGILK was detected in wild-type *S. cerevisiae*, in both glucose- and galactose-supplemented media (**Figure 3A**). The Hc-LPT1-specific peptide LTAFCWNVHDGR was detected only in complemented strains that were grown in galactose-supplemented medium (**Figure 3A**). The abundance of the peptide KGEELEIVGHNSTPLK from the house keeping protein elongation factor Tu (TUF1) was similar across different samples (**Figure 3A**).

**Figure 3.**
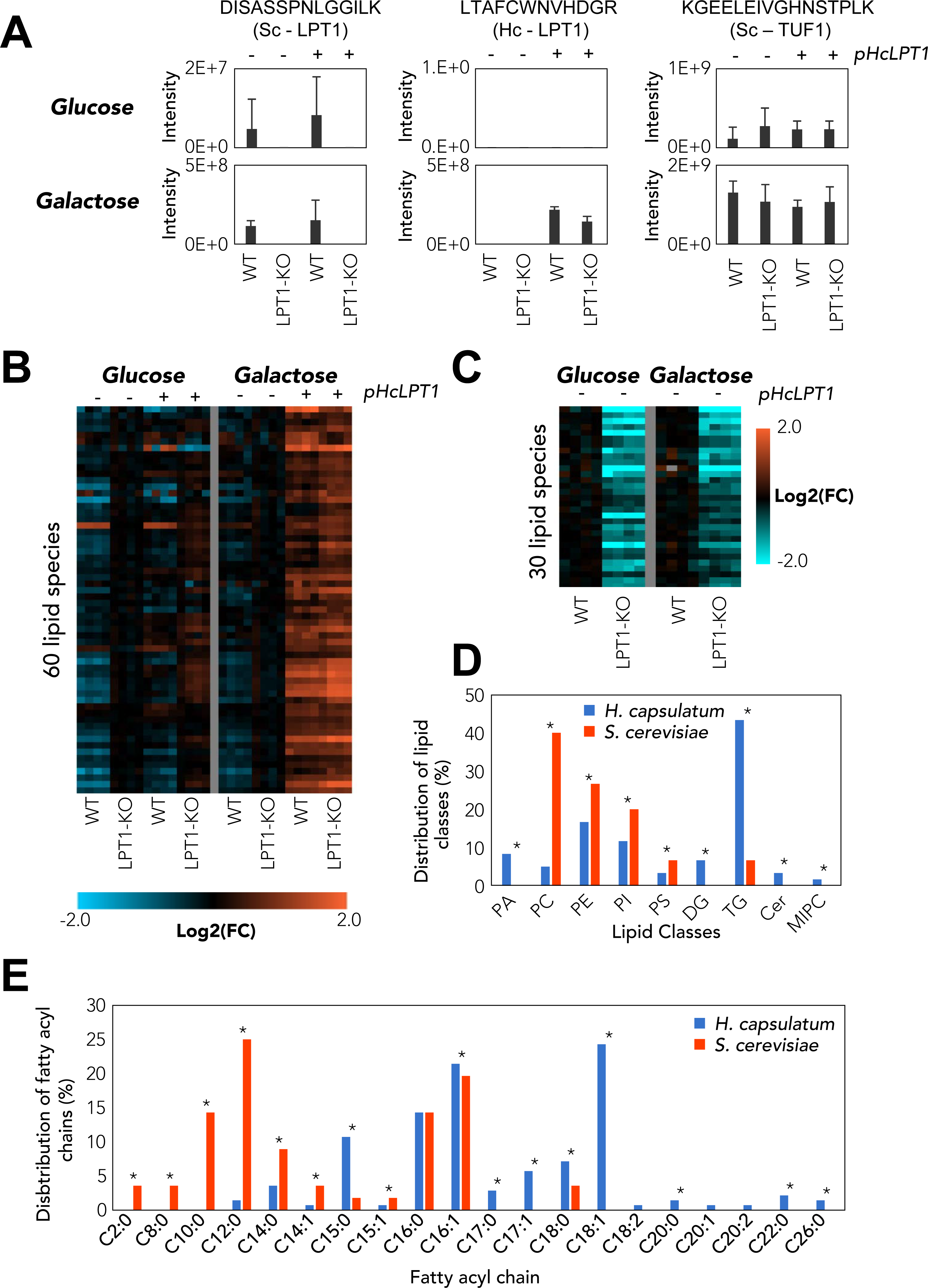
Analysis of *Saccharomyces cerevisiae* (Sc) and *Histoplasma capsulatum* (Hc) lysophospholipid acetyltransferase LPT1 genes. The analysis was done in wild-type and LPT1-knockout (KO) (*Lpt1* -/-) *S. cerevisiae* strains complemented or not with the pESC-URA plasmid with Hc-LPT1 gene under galactose-inducible promoter. *S. cerevisiae* WT and LPT1-knockout (KO) strains were grown in YNB medium supplemented with glucose (Glc) or galactose (Gal). (A) Protein abundance of Sc and Hc LPT1 proteins measured by the intensity of specific peptides in the LC-MS/MS analysis. Elongation factor Tu (TUF1) was used as a loading control. As expected, Sc LPT1 was only detected in the WT strain, whereas Hc LPT1 was detected only in strains transformed with the plasmid and induced with galactose. (B-C) Heatmaps of identified HcLPT1 (B) and ScLPT1 (C) products (p≤0.05 compared the complemented vs control strains (B) or LPT1-KO vs. WT strains (C). The experiments were done in two independent batches delimited by the vertical grey lines. (D-E) Distribution of the lipid classes (D) and fatty acyl groups (E) of the HcLPT1 and ScLPT1 products. *p≤0.05 by Fisher’s exact test.

The lipidomics analysis showed that Hc-LPT1 complementation increased the abundances of 60 lipids, including phosphatidic acid (PA), PC, PE, phosphatidylinositol (PI), PS, DG, TG, Cer and mannosyl-inositolphosphoceramide (MIPC) (**Figure 3B-D**, **Table S11**). The Hc-LPT1 complementation also significantly reduced the levels of 77 lipids, including 5 LPC, 3 lysophosphatidylethanolamines (LPE), 2 sphinganines and 3 DG (**Table S12**). On the other hand, disruption of the Sc-LPT1 gene reduced the levels of 30 potential products, including PC, PE, PI, PS and TG. Notably, the proportion of acylated products between the LPT1 from the two species was very different, with PC, PE, PI and PS being more acylated by the *S. cerevisiae* homolog and PA, DG, TG, Cer and MIPC by the *H. capsulatum* counterpart (**Figure 3D**, **Table S13**). The Sc-LPT1 showed a preference for production of lipid species containing short fatty acyl chains, with all the identified products having chains with ≤ 14 carbons (**Figure 3E**). Complementation with *Hc-Lpt1* gene showed a different phenotype, as this homolog produced lipid species containing odd-carbon number (C15 and C17) and longer (≥C18) fatty acyl chains (**Figure 3E**). Overall, the map provides a global view of lipids being produced by *H. capsulatum* along with catalytic enzymes.

### Evolutionary divergent lipid metabolic pathways as drug targets

We next focused on pathways that could be targeted for antifungal chemotherapies. The ideal drug target is essential for the fungal survival and divergent (substantially different or absent) compared to humans to allow specificity and reduce the chances of side effects. Therefore, we searched the lipid metabolic map for lipid pathways with enzymes that were present in *H. capsulatum*, but had no homologs in humans or *S. cerevisiae* as potential candidates for anti-*H. capsulatum* drug targets. The most divergent pathways were sphingolipid and fatty acid metabolism (**Figure 4A-B**). *S. cerevisiae* mainly produces (glyco)inositolphosphoceramides and humans produce complex glycosphingolipids, whereas *H. capsulatum* produces both (glyco)inositolphosphoceramides and simpler glycosphingolipids (**Figure RE 4A**).

**Figure 4.**
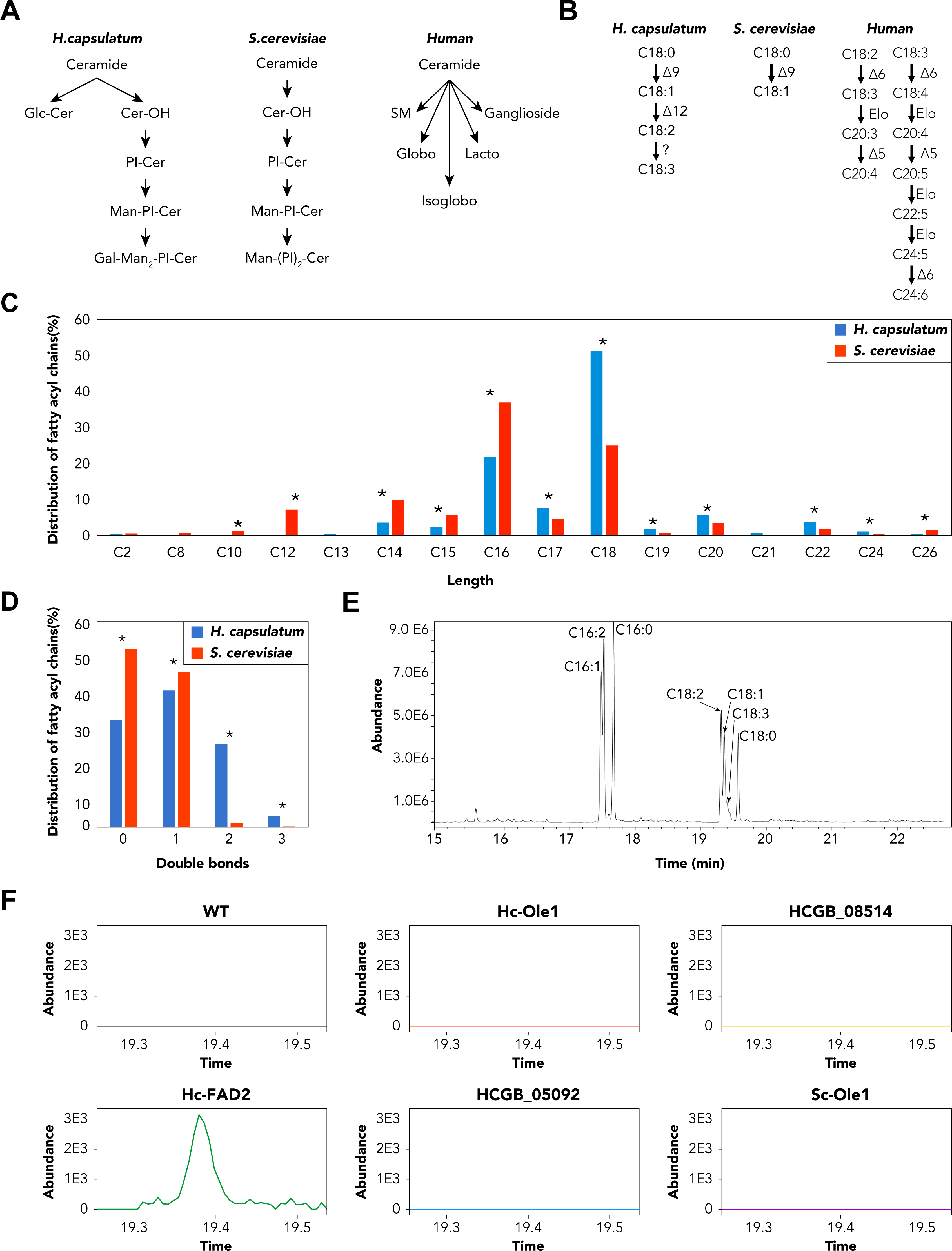
Sphingolipid and fatty acid metabolism in *Histoplasma capsulatum*. (A) Sphingolipids of *H. capsulatum*, *Saccharomyces cerevisiae* and humans. (B) Fatty acid desaturase pathways in *H. capsulatum*, *S. cerevisiae* and humans. (C) Distribution of carbon chain lengths of fatty acyl groups from lipids in global lipidomic analysis. (D) Distribution of double bonds on fatty acyl groups from lipids in global lipidomic analysis. (E) Representative chromatogram of fatty acid analysis from *S. cerevisiae* cells expressing *H. capsulatum* FAD2 gene. The analysis was done in *S. cerevisiae* strain transformed with the pESC-URA plasmid with Hc-FAD2 gene under galactose-inducible promoter. Cells were grown in YNB medium supplemented with galactose. The chromatogram shows the production of fatty acids with 2 and 3 double bonds. (F) Extracted ion chromatogram of α-linolenic acid (C18:3 – 9Z, 12Z, 15Z) of *S. cerevisiae* cells expressing various fatty acid desaturase candidates. α-linolenic acid was only detected in cells expressing the *H. capsulatum* FAD2 gene. Statistical test: *p≤0.05 by Fisher’s exact test.

**Figure 5.**
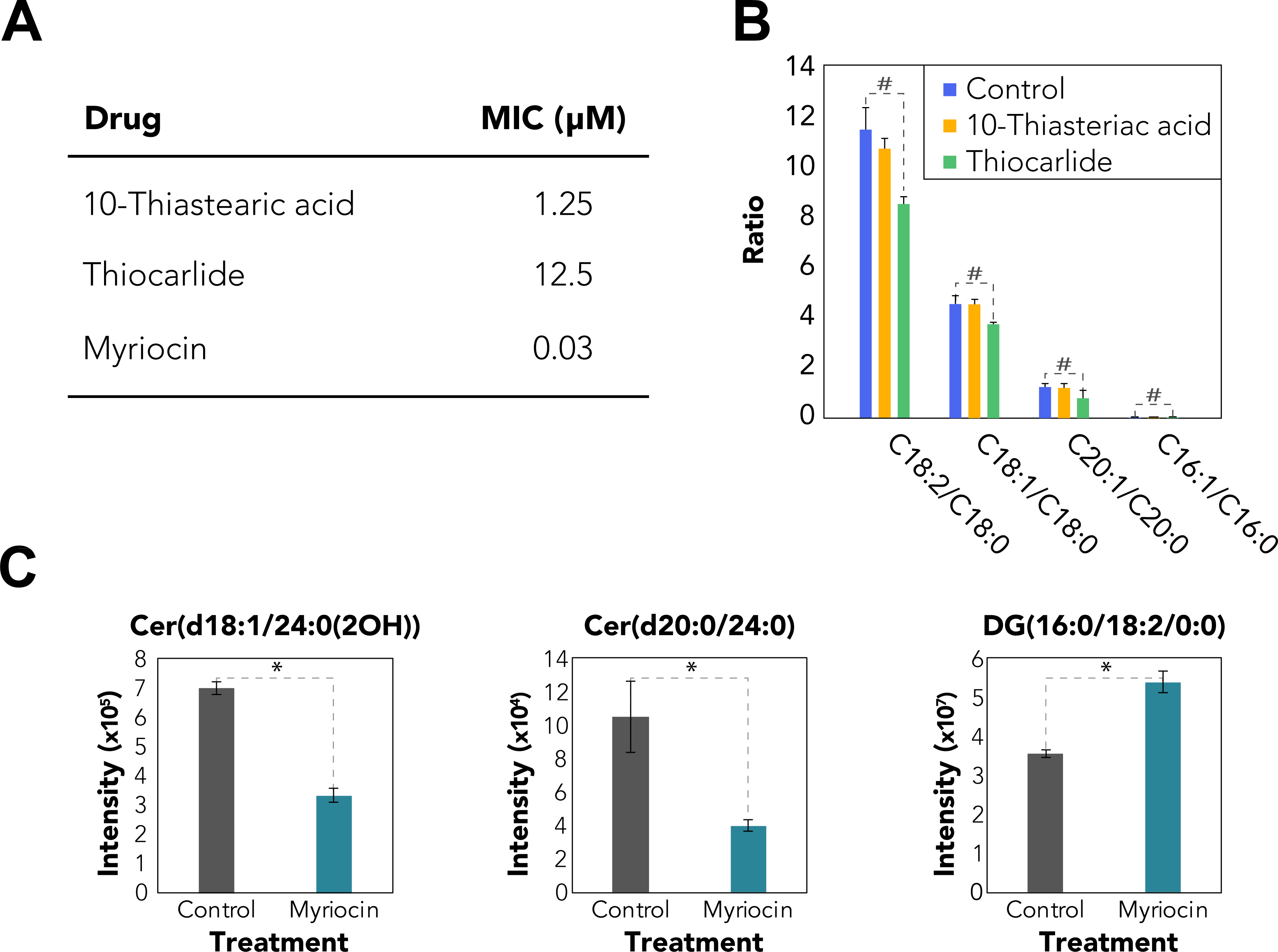
Targeting sphingolipid and fatty acid desaturation pathways for drug development. (A) Minimum inhibitory concentration (MIC) of 10-thiastearic acid, thiocarlide and myriocin. The MICs were obtained from 5 independent experiments with 4 replicates each. (B) Effect of fatty acid desaturase inhibitors in the fatty acid profile. Cells were grown in one half of each compound MIC and submitted to fatty acid analysis. The values represent the ratio of peak areas of unsaturated/saturate fatty acids. (C) Effect of myriocin on the lipidomics profile. Cells were grown in one half of myriocin MIC and submitted to fatty acid analysis. The values represent the mass spectral signal intensity on the apex of the peak. *p≤0.05 by Student *t*-test when comparing Control and treatments.

In fatty acid metabolism, humans are unable to perform the first two steps of fatty acid desaturation and instead uptake the respective unsaturated fatty acids from food, while *S. cerevisiae* has only one fatty acid desaturase gene (delta-9 desaturase, also known as oleate synthase *Ole1*) in its genome (**Figure 4B**). In addition to Ole1, *H. capsulatum* has a gene annotated as delta-12 desaturase (*FAD2*) and two uncharacterized desaturases (HCBG_08514 and HCBG_05092), which collectively might be responsible for the production of fatty acids with 2 and 3 double bonds (**Figure 4B**). We examined the distribution of fatty acyl chains that are incorporated into *H. capsulatum* and *S. cerevisiae* lipids. The results showed that both fungi incorporate fatty acyl groups with different chain lengths into their lipids. *S. cerevisiae* incorporates significantly more fatty acyl chains with less than 16 carbons (by Fisher’s exact test), whereas *H. capsulatum* has significantly more lipids with fatty acids longer than 17 carbons (**Figure 4C**). In terms of unsaturation, *S. cerevisiae* had almost exclusively saturated lipids or fatty acyl chains with 1 double bound, whereas *H. capsulatum* had fatty acyl chains with 2 or 3 double bonds (**Figure 4D**). To develop further insights on the enzymes responsible for desaturating these fatty acids, we expressed all 4 *H. capsulatum* desaturase candidates in *S. cerevisiae* by galactose induction, and determined the fatty acid composition by methylating them and analyzing them by GC-MS. We also included *S. cerevisiae* Ole1 as a positive control for over-expression. *S. cerevisiae* Ole1 over-expression increased the ratios of C14:1/C14:0, C16:1/C16:0 and C18:1/C18:0 by 4.0, 3.0 and 9.4 fold, respectively (**Figure S3A**), whereas the *H. capsulatum* Ole1 increased the ratios of C14:1/C14:0, C16:1/C16:0 and C18:1/C18:0 by 11%, 24% and 81%, respectively (**Figure S3A**). The expression of HCBG_08514 and HCBG_05092 had almost no impact on the saturation levels of the fatty acids (**Figure S3A**), which could be due to the lack of the precursors in *S. cerevisiae*. C16:2 (9Z,12Z-hexadecadienoic acid), C18:2 (9Z,12Z – linoleic acid) and C18:3 (9Z,12Z,15Z – α-linolenic acid) were only detected when *H. capsulatum* FAD2 was expressed (**Figures S3A-B**). A composition analysis of *H. capsulatum* fatty acids revealed two low abundance peaks of C18:3, one being α-linolenic acid and a second peak that did not match to any available standard (**Figures 4E-F** **and S3C**). These results showed that *H. capsulatum* FAD2 is indeed a bifunctional delta-12/delta-15 desaturase, which helped to further curate the lipid metabolic map (**Figure 2**, top left). Similar to sphingolipids, the *H. capsulatum* fatty acid desaturase pathway represents a potential target.

To perform a proof-of-concept that sphingolipid and fatty acid desaturation pathways can be targeted for drug development, we tested the effects of myriocin (serine-palmitoyltransferase inhibitor, which blocks the first step of sphingolipid biosynthesis pathway), and 10-thiastearic acid and thiocarlide (two fatty acid desaturase inhibitors) on *H. capsulatum* growth. Myriocin, 10-thiastearic acid and thiocarlide showed minimum-inhibitory concentrations (MICs) of 0.03, 1.25 and 12.5 μM, respectively (**Figure 5A**). To confirm that these compounds target fatty acid desaturases and sphingolipid biosynthesis, we performed a fatty acid analysis on treated and unexposed yeast cells. Cells were grown for 48 hours with one half of the MIC and fatty acids were extracted, methylated and analyzed by GC-MS. 10-Thiastearic acid did not affect the ratio between unsaturated and saturated fatty acids (**Figure 5B**), suggesting that this compound might target other pathways in *H. capsulatum* rather than the fatty acid desaturation. As expected, thiocarlide reduced the C18:2/C18:0 peak area ratio from 11.4 to 8.5 (25% reduction), the C18:1/C18:0 ratio from 4.5 to 3.7 (18% reduction) and the C20:1/C20:0 ratio from 1.3 to 0.8 (39% reduction) (**Figure 5B**). Conversely, the C16:1/C16:0 ratio had a slight increase from 0.036 to 0.044 (22% increase) (**Figure 5B**). We also performed a lipidomic analysis on cells grown with or without one half of myriocin MIC for 48 hours. As expected, the most abundant ceramide species, Cer(d18:1/24:0(2OH)) and Cer(d20:0/24:0), were reduced by 52% and 62%, respectively (**Figure 5C**). As a control of a lipid from an unrelated class, the level of the most abundant DG, DG(16:0/18:2/0:0), increased by 51% (**Figure 5C**).

To further test the fatty acid desaturation and sphingolipid pathway as anti-*H. capsulatum* drug targets, we tested the ability of these inhibitors to reduce intracellular infection. We chose alveolar macrophages as they are the primary cells targeted by *H. capsulatum* in infection (30). We performed a toxicity assay with concentrations up to 64-fold higher than the MICs. Myriocin 10-thiastearic acid did not affect the viability of AMJ2 alveolar macrophages, while thiocarlide had a small effect, reducing the viability of the cells by approximately 30% on the highest tested concentrations (**Figure 6A**). We tested a low (2x MIC: 0.06 μM myriocin, 2.5 μM 10-thiastearic acid and 25 μM thiocarlide) and a high concentration (2 μM myriocin, 80 μM 10-thiastearic acid and 50 μM thiocarlide) to determine the ability of the compounds to reduce intracellular infection. In low concentrations, myriocin, 10-thiastearic acid and thiocarlide reduced intracellular *H. capsulatum* load by 21.4%, 19.2% and 30.1%, respectively (**Figure 6B**), while the higher concentration further reduced this by 32.0%, 21.4% and 37.4% (**Figure 6C**). These results validate the fatty acid desaturation and sphingolipid pathways as a potential target for developing anti-*H. capsulatum* drugs, and that thiocarlide can reduce intracellular fungal load in alveolar macrophages.

**Figure 6.**
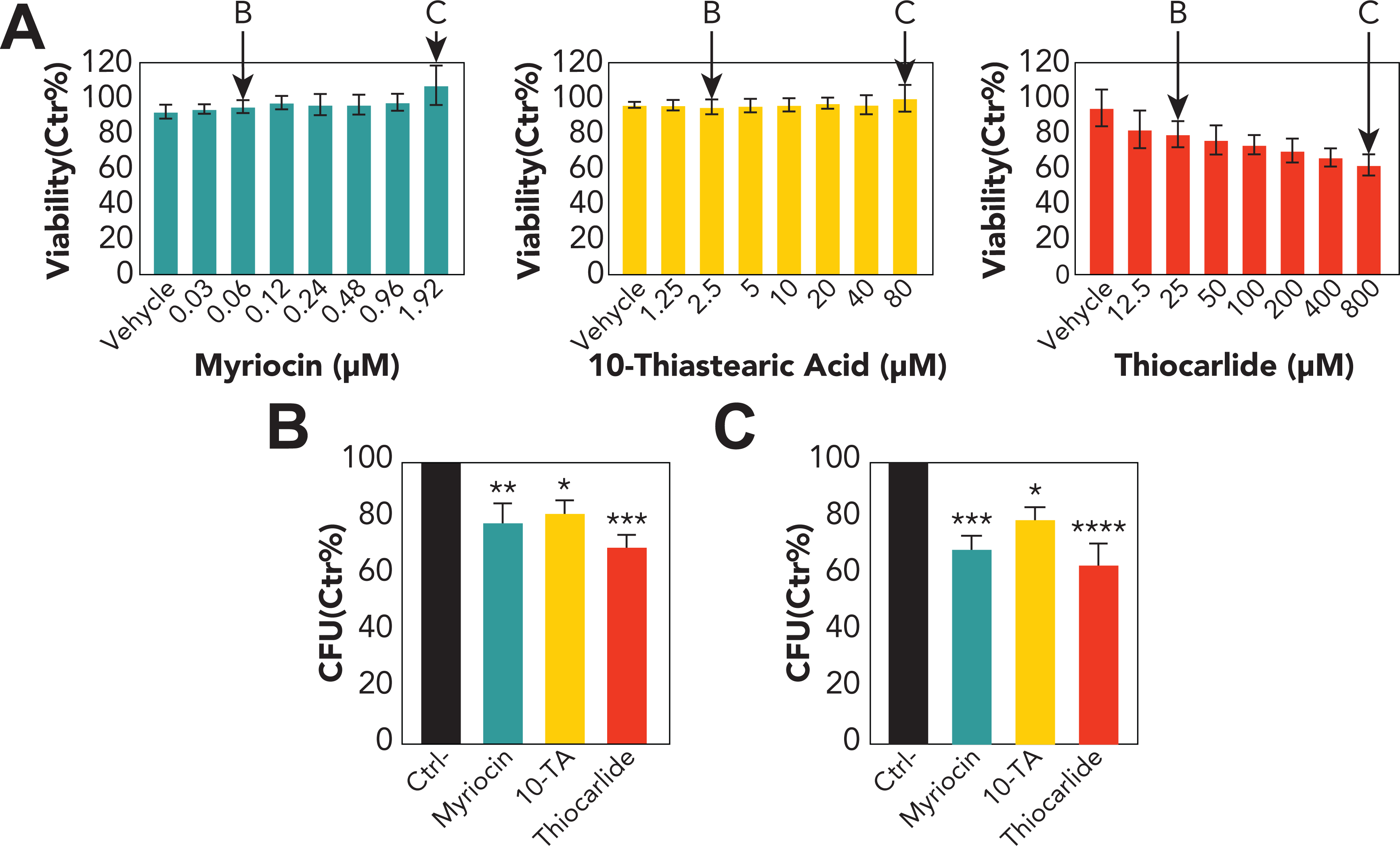
Toxicity and efficacy of lipid metabolism inhibitors against intracellular infection of AMJ2 alveolar macrophages with *Histoplasma capsulatum*. (A) Viability of AMJ2 alveolar macrophages, measured by MTT (3-(4,5-dimethylthiazol-2-yl)-2,5-diphenyltetrazolium bromide) assay, after exposed to different concentrations of myriocin, 10-thiastearic acid and thiocarlide. (B-C) Intracellular killing of *H. capsulatum* by AMJ2 alveolar macrophages treated with the low and high concentrations of myriocin, 10-thiastearic acid (10-TA) and thiocarlide. The concentrations are highlighted in panel A. Bars represent the combination of 4 independent experiments with 3 replicates each (n=12). *p≤0.05, **p≤0.01 and ***p≤0.001 by One-way-Anova followed by Dunnett’s test comparing control and treatments.

## DISCUSSION

Here we developed a global map of the *H. capsulatum* lipid metabolism by incorporating genomic, proteomic and lipidomic information, including relative abundances of proteins and lipids. We found accordance of the protein abundance levels with their functions in the lipid metabolism. For instance, the proteins involved in the synthesis of the major lipid components--the fatty acids and sterols--were enriched among the high and moderate abundant proteins, whereas the proteins related to the synthesis of specific lipid head groups were enriched in the low to very low abundant proteins. This might be a consequence of cell resource optimization as protein synthesis is one of the most energetically expensive tasks in cells (31). Our lipid map also showed that TG, PC and PE are the most diverse classes of lipids, which is in agreement with the fact that they are most abundant ones (32). Our results show that the diversity of TG, PC and PE species could be due to fatty acid remodeling, as these lipids are products of the LPT1 acyltransferase. LPT1 transfers acyl groups to a variety of fatty acyl chains, including odd-carbon chains (C15 and C17) and long-chain (≥C18) fatty acids. Our experiments with LPT1 also helped to curate the lipid map since the *S. cerevisiae* homolog has a different specificity for substrates, that being a major impact in producing phospholipids with short fatty acyl chains. In *S. cerevisiae*, the substrates of LPT1 have been shown *in vitro*, using radioactive precursors to acylated LPC, LPG, LPA, LPE, LPI and LPS (27). The acylation of LPA, a central precursor for all glycerolipids and glycerophospholipids, suggests that the LPT1 impact could be indirect. Therefore, further investigation will be needed to determine if this enzyme can directly acylate DG into TG. We also curated the fatty acid desaturation pathway by expressing *H. capsulatum* desaturase candidate genes in *S. cerevisiae*. We validated that Ole1 has delta-9 desaturase activity. In addition, we show that FAD2 is indeed a bifunctional delta-12/delta-15 desaturase. In fungi, delta-15 and bifunctional delta-12/delta-15 desaturases evolved from delta-12 desaturases in multiple independent gene duplications, which complicates the assignment of their specificity based on their sequence (28). This scenario is a little different in *H. capsulatum*, as *H. capsulatum* FAD2 evolved to have both activities within a single gene copy rather than duplicating the gene.

We identified the fatty acid desaturation and the sphingolipid pathways as divergent points in the lipid metabolism pathways of *H. capsulatum* vs. *S. cerevisiae* and humans. We showed that both fatty acid and sphingolipid pathways are potential targets for developing anti-histoplasmosis drugs. Thiocarlide (also known as isoxyl) is a potent inhibitor (nanomolar range) of *Mycobacterium tuberculosis* delta-9 stearoyl desaturase and was used as a second line of anti-tuberculosis drugs in the 1960s (33,34). However, despite its low toxicity, thiocarlide was not as effective during clinical trials, so is no longer used to treat patients (34-36). One possible explanation for the clinical trial failure is the poor solubility of the compound in water. Since then, other drug delivery vehicles have been tested with promising results *in vitro* (37). Delta-12 oleate desaturase has been validated as a drug target in the protozoan parasite, *Trypanosoma cruzi* (38). Indeed, thiocarlide inhibited *T. cruzi* growth in low micromolar range and reduced the levels of unsaturated fatty acids by approximately 30% (at 10 μM concentration) (39). These numbers are similar to the MIC and reduction in unsaturated fatty acid level for *H. capsulatum* (**Figure 5A-B**). The small changes in unsaturated fatty acid levels were associated with a drastic reduction in cell viability, which suggests that maintenance of homeostasis in membrane fluidity is crucial for life. Moreover, thiocarlide was able to reduce intracellular infection with *H. capsulatum*. The small reduction in intracellular infection might be due to limited diffusion through the host cell membrane, which is observed in current antifungal drugs such as voriconazole, and increased concentrations are required to kill intracellular fungi (40). Such desaturase inhibitors should also be tested in other pathogenic fungi, as many of them produce polyunsaturated fatty acids (41,42).

Our data also showed that 10-thiastearic acid might target a different pathway rather than fatty acid desaturation. This is expected to some extent since 10-thiastearic acid is a fatty acid analog; therefore, it is more likely to inhibit other processes of fatty acid metabolism. However, this type of molecule should still be considered as a drug candidate due to its antifungal activity and potential low toxicity to humans. Our data showed no toxicity in AMJ2 alveolar macrophages with up to 64-folds the MIC of 10-thiastearic acid. In addition, the 10-thiastearic acid analog tetradecylthioacetic acid has a low toxicity in humans up to 1 g/day (43). Engineered delivery methods, such as liposomes, and structural modifications, such as esterification, can further improve the bioactive compound to reach the intracellular milieu. Therefore, 10-thiastearate might still be useful for treating histoplasmosis but further development still needs to be performed to test their efficacy.

We also performed a proof-of-concept that sphingolipids are promising targets for anti-histoplasma drug development by inhibiting the first step of the sphingolipid pathway with myriocin. Myriocin kills *Candida albicans, C. auris* and *Aspergillus fumigatus* in nanomolar to micromolar concentrations (17,44,45). *H. capsulatum* is more sensitive to myriocin as its MIC was 30 nM (**Figure 5B**). Myriocin was also able to reduce the intracellular fungal load. Unfortunately, myriocin has immunosuppressant activity (46,47), but this has been explored for simultaneously killing the fungus and reducing the pathogenic inflammation of the lungs in cystic fibrosis mice infected with *A. fumigatus* (48). The fact the *de novo* sphingolipid synthesis is essential for *H. capsulatum* opens the opportunity to explore other inhibitors of serine palmitoyltransferase, or other enzymes in the pathway. Indeed, other enzymes of the fungal sphingolipid pathway have been validated as drug targets. In *C. neoformans*, acylhydrazones have been shown to be potent inhibitors of the glucosylceramide synthesis (nanomolar range) and excellent antifungal drug candidates (13,49). Sphingolipids have also been shown to be a mechanism of azole-resistance in *C. albicans* (50) and at the same time, excellent targets for synergistic drugs across multiple species (51).

In conclusion, we built a detailed map of *H. capsulatum* lipid metabolism based on omics data. Our data identified defined regions of the *H. capsulatum* lipid metabolic pathway that can be targeted for drug development. The lipid metabolic map is a valuable resource to the community and its use can help in the discovery of other functions of lipids in fungal physiology and pathogenesis.

### Limitations of study

One limitation of this study is that *H. capsulatum* is a fungus with few genetic engineering tools, which makes the execution of gene knockout or knockdown experiments highly challenging. Therefore, we utilized the gene complementation system in *S. cerevisiae* to study the specificity of genes from the lipid metabolism. We found dozens of *H. capsulatum* LPT1 products and determined the double bond positions catalyzed by FAD2, but the complementation system can generate false negatives due to the absence of specific substrates in *S. cerevisiae*. The different lifestyles between *H. capsulatum* and *S. cerevisiae* limits some of the comparisons between the lipid composition of these two organisms. Another limitation of our study is that the *H. capsulatum* lipidomic analysis was performed in a single growth condition, which might not fully recapitulate the *in vivo* environment during infection. Also, we have previously shown that different growth conditions have an impact on the composition of extracellular vesicles, including variations in lipid profiles (52). Therefore, the analysis of different growth conditions and forms of the fungus might lead to different lipid profiles and even the identification of additional lipid species. The current work was performed in a single *H. capsulatum* strain (G217B). Strain-dependent variations regarding lipid and protein profiles are expected among different fungal strains of the same species, which needs to be further studied in future work.

## Material and Methods

### Cells

*Histoplasma capsulatum* G217B strain and murine alveolar macrophages AMJ2 was purchased from the American Type and Culture Collection (ATCC, Manassas, VA). *S. cerevisiae* deficient LPT1 gene (YOR175C knockout (KanMX), strain 12431) and wild-type strains (BY4742; mat alpha, his3Δ1 leu2Δ0 lys2Δ0 ura3Δ0) were purchased from Dharmacon (Lafayette, CO).

### *Saccharomyces cerevisiae* transformation

Strains were maintained on YPD (2% glucose, 10 g/L peptone, 10 g/L yeast extract) or YNB with amino acid supplements (2% glucose, 1.7 g/L YNB salts without ammonium sulfate or amino acids (Difco), 5 g/L ammonium sulfate (Fisher), +/- 50 mg/L leucine, lysine, histidine (LLH) and uracil). The strains were transformed by a lithium-acetate-heat shock method (53) with a plasmid containing the *H. capsulatum* acyltransferase LPT1 (Uniprot ID C0NZS2), or desaturases Ole1 (Uniprot ID C0NLE5), FAD2 (Uniprot ID C0NKL1), HCBG_08514 (Uniprot ID C0NZD4) and HCBG_05092 (Uniprot ID C0NPL2) genes and *S. cerevisiae* Ole1 (Uniprot ID P21147) that were codon optimized, synthesized and cloned into the Gal-inducible plasmid pESC-URA by GenScript (Piscataway, NJ). Briefly, 500 μL of cells grown in YPD for 16 h were spun down, and the supernatant removed. Added to the BY4742 and #12431 cells were: 5 μg of herring sperm DNA (5 μL of 10 mg/mL stock), and 400 ng of GenScript plasmid DNA, 500 μL of PLATE buffer (40% PEG 3350, 0.1 M lithium acetate, 10 mM Tris HCl (pH 8.0), 1 mM EDTA), and 57 μL DMSO. Cells were incubated at room temperature for 15 min, followed by 15 min at 42 °C. Cells were collected by brief centrifugation of 5 s at 8000 x g, supernatant removed, resuspended in PBS, then plated on YNB-2% glucose-LLH 2 % agar plates without uracil. Resulting colonies were screened for plasmid presence by PCR of a portion of the pESC-URA-lpt1 plasmid (Forward primer: 5’-TTGGAAACAGCTCCAAATCC-3’, Reverse primer: 5’ CCCAAAACCTTCTCAAGCAA-3’; ordered from ThermoFisher Oligos) and preserved as glycerol stocks.

### *S. cerevisiae* plasmid expression assays

For the Hc_LPT1 and desaturase expression assays, cells were pre-cultured overnight in YNB 2% glucose LLH +/- uracil to accommodate the non-plasmid strain auxotrophy as a negative control. For biomass collection during induction, which often slows or prevents cell division, cells were inoculated to an OD of 0.5 into YNB-2% galactose LLH +/- uracil, and collected after induction of plasmid expression for 17 hours. LPT1 cultures were also analyzed after 17 hours grown in glucose LLH +/- uracil (uninduced). Cultures were collected by spinning 25 mL of culture in a tabletop centrifuge (VWR symphony 4417 R) at 2000 x g for 2 min with a swinging bucket rotor fitting 50-mL conical tubes. Pellets were collected into pre-weighted tubes and washed 2X with TBS before flash freezing in liquid nitrogen. All samples were done in 4 replicates as input for downstream omics analysis.

### *Histoplasma capsulatum* cell culture

Yeasts of *H. capsulatum* G217B were grown in Ham’s F12 medium at 37°C under constant shaking, and used 3 days after its inoculation, as previously described (54). Cells were harvested by centrifuging at 2000 x *g* for 2 min and washed twice with PBS. All the sample preparation and analysis were done in 4 replicates and in a randomized order to ensure the biological significance and prevent batch effects.

### Sample extraction and fractionation on Silica 60 column

Samples were extracted using two different approaches. For global proteomics and lipidomics analyses, samples were submitted to <u>M</u>etabolite, <u>P</u>rotein and <u>L</u>ipid <u>Ex</u>traction (MPLEx) as previously described (23). For the fractionation study, cells were extracted twice with chloroform:methanol (2:1, v:v) and twice chloroform:methanol:water (1:2:0.8, v:v:v)) as described elsewhere (19). Extracted lipids were fractionated into neutral lipids, fatty acids and phospholipid fractions using Silica 60 columns (55) and dried in a vacuum centrifuge.

### Proteomic analysis

Extracted proteins were digested and analyzed by LC-MS/MS as described (56). Data were analyzed with MaxQuant software (v.1.5.5.1) by searching against the *H. capsulatum* G186AR (downloaded August 15, 2016) and *S. cerevisiae* S288c (downloaded January 11, 2018) sequences from Uniprot Knowlegdgebase, considering protein N-terminal acetylation and oxidation of methionine as variable modifications, and cysteine carbamidomethylation as a fixed modification. The remaining parameters were set as the software default. Protein abundances were estimated by the intensity-based absolute quantification (iBAQ) method (57). The intensities were normalized by the total iBAQ sum of each sample and expressed as relative number of protein copies (percentage from total). Proteins were classified according to their abundance converting number of copies described by Beck et al. (24) to the relative number of protein copies (%). Function-enrichment analysis was performed as described previously (58).

### Lipid analysis by liquid chromatography-tandem mass spectrometry

Total lipid extracts and phospholipid fraction from the Silica 60 column were resuspended in methanol and subjected to LC-MS/MS analysis as previously described (59). To assess the technical variability, we spiked in the SPLASH Lipidomix standard (Avanti Polar), which contains a mix isotopically labeled lipids. Lipid species were identified using LIQUID, which matches spectra against comprehensive database of lipids species, including all the species contained in LipidMaps in addition to several other classes of lipids recently described in the literature (60). Identified species were manually inspected for validation based on isotopic distribution, and head group and fatty acyl fragments. The features of the identified lipid species were extracted and aligned using MZmine (61). For comparative purposes, lipids were considered significantly different with p ≤ 0.05 by *t*-test considering equal variance and unpaired samples. The distribution of fatty acyl groups was done by counting individual fatty acyl groups and were considered significantly different with a p ≤ 0.05 by Fisher’s exact test.

### Fatty acid and sterol analyses

Fatty acid were methylated with anhydrous methanolic HCl (1.2 N) for 1 h at 100°C, and extracted by adding equal volumes of water and hexane. Sterols were treated with 30 mg/mL methoxyamine in pyridine for 90 min at 37 °C with shaking, and derivatized with N-methyl-N-(trimethylsilyl)trifluoroacetamide (MSTFA) (Sigma-Aldrich) with 1% trimethylchlorosilane (TMCS) (Sigma-Aldrich) at 37°C with shaking for 30 min (62). Derivatized FAMEs and sterols were then analyzed in an Agilent GC 7890A using an HP-5MS column (30 m × 0.25 mm × 0.25 μm; Agilent Technologies, Santa Clara, CA) coupled with a single quadrupole MSD 5975C (Agilent Technologies). Samples were injected (splitless) into the port set at 250°C with an initial oven temperature of 60°C. After 1 minute the temperature was increased to 325°C at a rate of 10°C/minute, and finished with a 5-minute hold at 325°C. The data files were calibrated with external calibration of FAME standards (C8-28, Sigma-Aldrich) and deconvoluted with Metabolite Detector as stated previously (63). Molecules were identified by library matching against the FiehnLib (64) with additional in-house entries, the Wiley Fatty Acids Methyl Esters Mass Spectral Database and the NIST17 GC-MS spectral library.

### Evaluation of lipid biosynthesis inhibitors on *H. capsulatum* axenic and intracellular growth

*H. capsulatum* yeasts were washed with PBS and suspended in Ham’s F12 and loaded into plates with serially diluted compounds at a final cellular density of 2.5 x 10^4^ cells/mL in 4 replicates. Cells were grown at 37 ⁰C under constant shaking for 7 days and minimal drug concentration with no visual growth was considered as MIC. To evaluate the effect of lipid synthesis inhibition of *H. capsulatum*, triplicates of 10^6^ cells/mL were incubated in Ham’s F12 at 37 ⁰C shaking at 250 rpm for 48 h with one half of the MIC of myriocin, 10-thiasteric acid and thiocarlide. Cells were washed with PBS and pellets were frozen for further lipid extraction and subsequent fatty acid and lipidomics analyses. To assess the ability of the compounds of killing intracellular fungi, murine alveolar macrophage cell line AMJ2 was plated onto 96-well plates and incubated overnight at 37 °C with 5% CO_2_. The macrophages were challenged with *H. capsulatum* (ratio of 2 yeasts per macrophage cell) in the presence or absence of myriocin, 10-thiasteric acid and thiocarlide for 48 h (see Figure s for concentrations). Macrophages were lysed with distilled water and the yeast suspensions were diluted and plated onto BHI-agar supplemented with 5% of sheep’s blood. Plates were incubated at 37 °C and colony forming units were counted after grown. To address cytotoxicity of the inhibitors upon alveolar macrophages, AMJ2 cells were plated (10^5^ cells/well) in 96-well plates, incubated overnight at 37 °C and 5% CO_2_. Cells were washed with PBS and incubated with serially diluted inhibitors. Solvent controls were made with the same solvent amount present in the highest concentration for each inhibitor. Plates were incubated for 24 hours at 37 °C and 5% CO_2_. MTT (3-(4,5-dimethylthiazol-2-yl)-2,5-diphenyltetrazolium bromide) was added to each well (50 µg/well) and incubated for 4 hours at 37 °C and 5% CO_2_. After the supernatant removal, the formazan crystals were solubilized with isopropanol and read with a plate reader at 570 nm. All values were normalized using the negative control (absence of drug or solvent).

### Construction of the *H. capsulatum* lipid metabolic map

We have built a preliminary G217B lipid map with a tool name VANTED v2.6.5 (65), using the *S. cerevisiae* model (25) as a starting point. Enzyme homologs were identified by blasting *S. cerevisiae* and *C. neoformans*, two of the best-characterized fungi in terms of lipid metabolism (25,66). We also based the map on the information available on the *H. capsulatum* from the lipid literature and lipids identified in the lipidomics analysis. The map was integrated by adding the abundance level of lipids using their relative mass spectrometry intensities within the same lipid class.

### Lipidomics and proteomics visualization tool

We developed a software tool based on the lipid metabolic map to automatically build a figure representing lipidomics and proteomics data. Colorblind-friendly palettes were applied to the relative abundance values and mapped to a static image displaying lipid metabolism pathways. The script was written in Perl using the GD graphics package.

## DATA AND SOFTWARE AVAILABILITY

Proteomics data were deposited into Pride repository (www.ebi.ac.uk/pride) under accession number PXD017734.

The code for the lipidomics visualization tool is available under the BSD 2-Clause License at GitHub under URL: https://github.com/wichne/LipidomicsMapTools

## AUTHOR CONTRIBUTIONS

D.Z.M., H.M.H., M.C.B., J.D.N and E.S.N. designed the research. D.Z.M. performed the *H. capsulatum* growth and drug testing. H.M.H., M.C.B., E.M.Z., J.E.K., K.K.W., E.L.B., X.Z., E.S.B. and K.J.B. performed the lipidomics and proteomics experiments. E.L.B. performed the genetic engineering of the *S. cerevisiae* strains. M.C.B., H.M.H., N.M.M. and Y.M.K. performed the gas chromatography-mass spectrometry analyses. E.S.N., G.C., J.D.Z., J.R.T. build the lipid metabolic map. W.C.N. developed the lipidomics/proteomics visualization tool. D.Z.M., H.M.H., M.C.B., S.P.C., X.Z., E.S.B, J.E.K., S.H.P., Y.M.K., M.R.G., J.D.N., and E.S.N. analyzed the data. M.R.G., E.S.B, S.H.P., J.D.N and E.S.N. contributed with reagents and resources. D.Z.M., H.M.H, M.C.B. and E.S.N. wrote the manuscript with inputs from the other authors. All the authors read and approved the final version of the manuscript.

## CONFLICTS OF INTEREST

The authors declare no financial conflicts of interest.

## Acknowledgments

The authors thank Drs. Igor Almeida, Rosa Maldonado, Milene Carmes Vallejo, Charles Ansong and Joshua Adkins for insightful discussions. Joshua D. Nosanchuk and Ernesto S. Nakayasu were supported by NIH R21 AI124797. Erin L. Bredeweg, Ernesto S. Nakayasu and Jennifer Kyle were supported by a Laboratory Directed Research and Development project from Pacific Northwest National Laboratory (PNNL). Erin S. Baker would also like to acknowledge support from her NIEHS P42 ES027704 award. Parts of this work were performed in the Environmental Molecular Science Laboratory, a U.S. Department of Energy (DOE) national scientific user facility at PNNL in Richland, WA.

---

**Figure S1.**
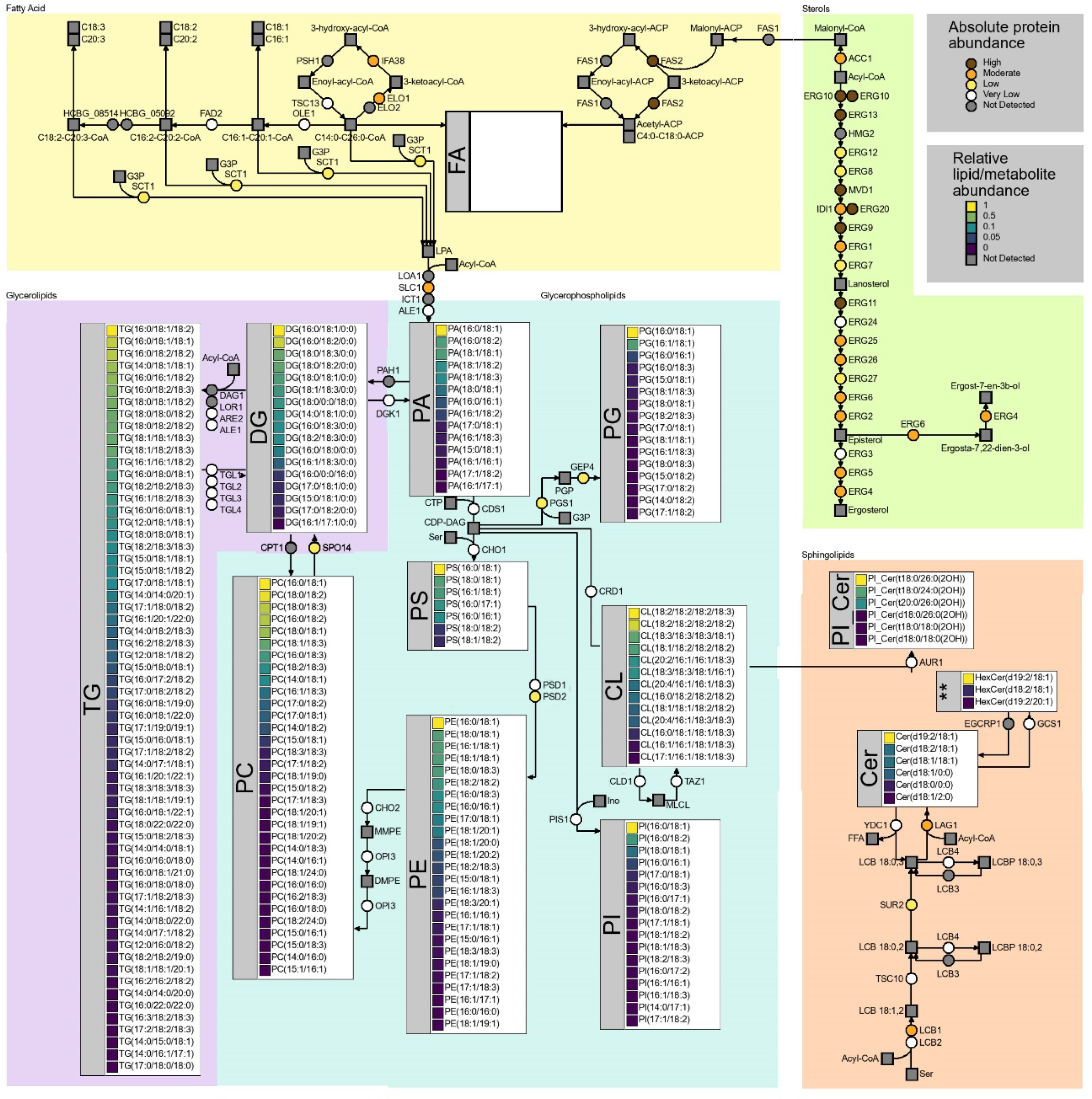
Lipid metabolic pathway visualization tool. The lipid metabolic pathway built for *Histoplasma capsulatum* was converted into a lipidomics and proteomics data visualization tool using a script written in Perl and using the GD graphics package. The tool generates color-coded expression values based on the omics measurements.

**Figure S2.**
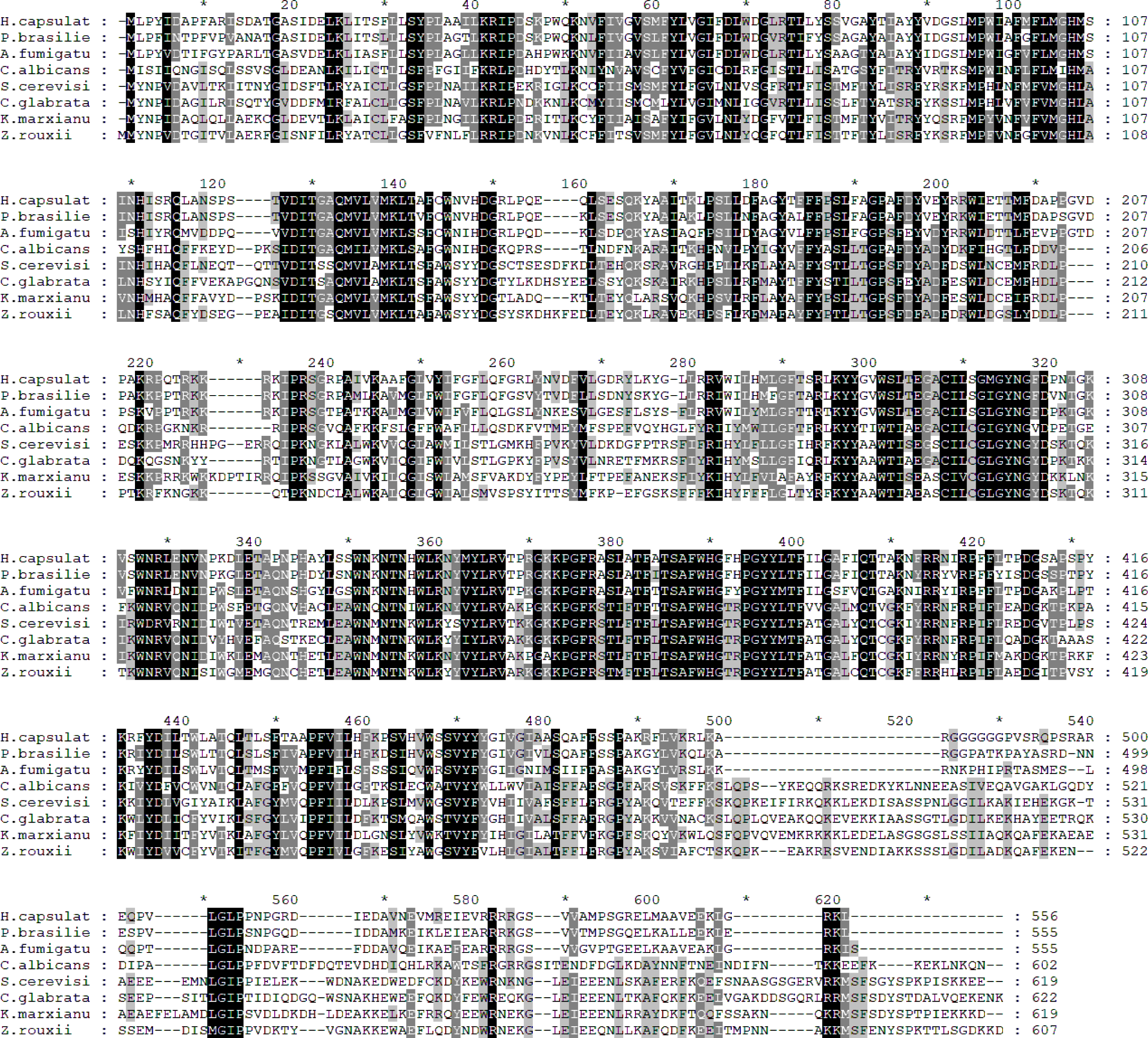
Sequence alignment of *Histoplasma capsulatum* lysophosphatidylcholine acyltransferase LPT1 candidate gene and various homologs from other fungal species. *H. capsulatum* MBOAT family protein (NCBI accession number: EEH03012) was aligned with sequences from *Paracoccidioides brasiliensis* Pb18 (89% similarity, NCBI accession number: EEH50158), *Aspergillus fumigatus* Af293 (81% similarity, NCBI accession number: EAL85553), *Candida albicans* P37039 (56% similarity, NCBI accession number: KHC88112), *Saccharomyces cerevisiae* S288C (53% similarity, NCBI accession number: DAA10947), *Candida glabrata* (52% similarity, NCBI accession number: KTB01140), *Kluyveromyces marxianus* DMKU3-1042 (53% similarity, NCBI accession number: BAO40203) and *Zygosaccharomyces rouxii* (53% similarity, NCBI accession number: GAV49200).

**Figure S3.**
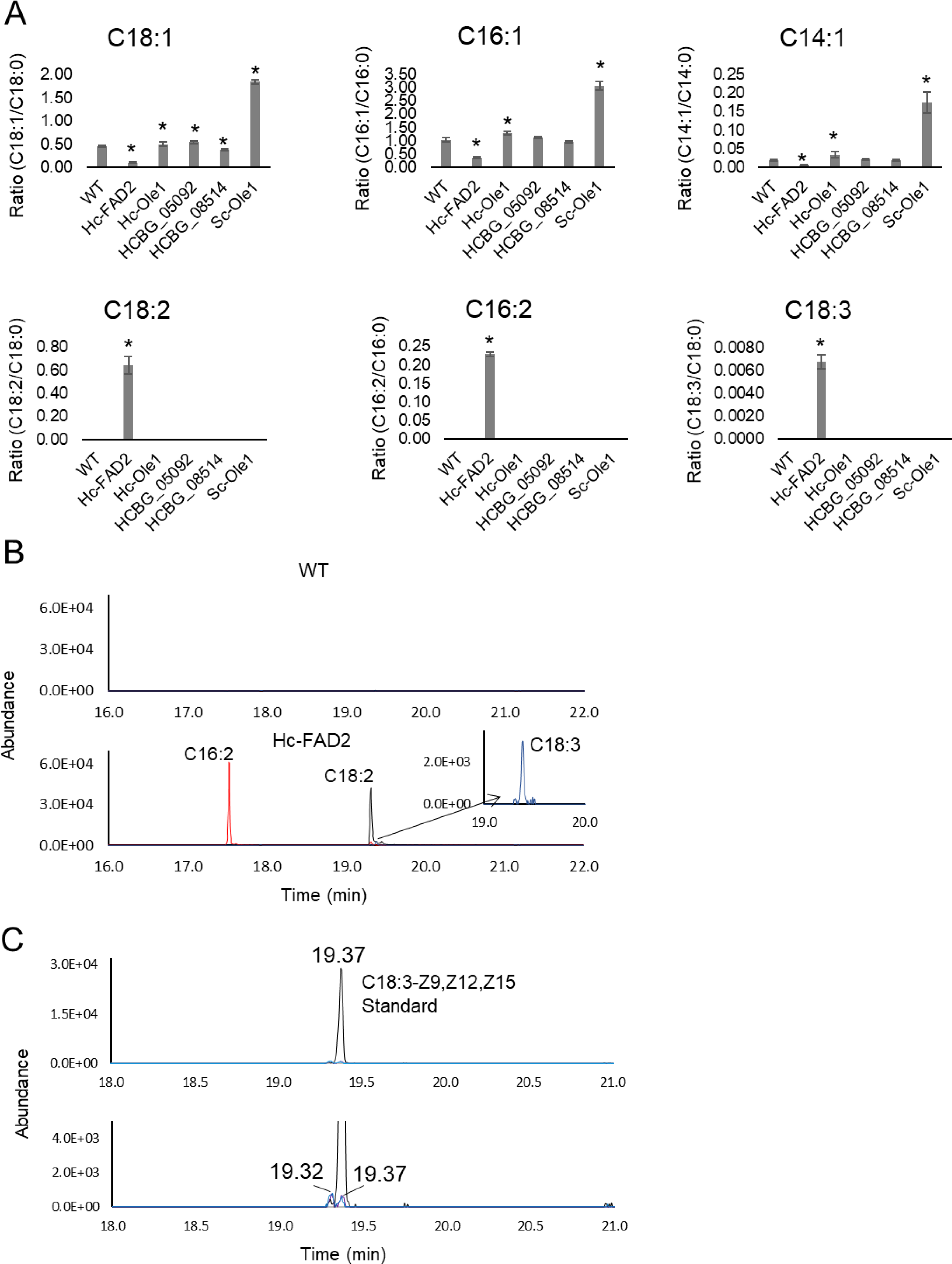
Fatty acid composition analysis. (A) The analysis was done in *S. cerevisiae* strain transformed with the pESC-URA plasmid with *H. capsulatum* Ole1 (Hc-Ole1), FAD2 (Hc-FAD2), HCBG_08514 and HCBG_05092 genes and *S. cerevisiae* Ole1 (Sc-Ole1) gene under galactose-inducible promoter. Cells were grown in YNB medium and supplemented with galactose for 17 h. Lipids were extracted and hydrolyzed, and the resulting fatty acids were methylated and analyzed by gas chromatography-mass spectrometry. Levels of unsaturated fatty acids were expressed as ratios compared to their saturated counterpart. The experiment was performed in biological 4 replicates. Statistical test: *p≤0.05 compared to the control using Student *t*-test. (B) Extracted ion chromatograms of fatty acids with 2 and 3 double bonds from control and *S. cerevisiae* strains expressing the *H. capsulatum* FAD2 gene. The chromatograms show that fatty acid species with 2 or 3 double bonds were not detected in wild-type *S. cerevisiae*. (C) C18:3 fatty acids presence in *H. capsulatum* lipids. Lipids were extracted and hydrolyzed, and the resulting fatty acids were methylated and analyzed by gas chromatography-mass spectrometry. Comparison with external standards identified the peak at 19.37 minutes as α-linolenic acid, while the peak at 19.32 remained unidentified.

## Notes

### Competing Interest Statement

The authors have declared no competing interest.

### Summary of Updates

We added new results on the toxicity of the tested compounds to alveolar macrophages and their ability to treat intracellular fungal infection.

## References

1. Bongomin, F., Gago, S., Oladele, R. O., and Denning, D. W. (2017) Global and Multi-National Prevalence of Fungal Diseases-Estimate Precision. J Fungi (Basel) 3

2. Chu, J. H., Feudtner, C., Heydon, K., Walsh, T. J., and Zaoutis, T. E. (2006) Hospitalizations for endemic mycoses: a population-based national study. Clin Infect Dis 42, 822–825

3. Armstrong, P. A., Jackson, B. R., Haselow, D., Fields, V., Ireland, M., Austin, C., Signs, K., Fialkowski, V., Patel, R., Ellis, P., Iwen, P. C., Pedati, C., Gibbons-Burgener, S., Anderson, J., Dobbs, T., Davidson, S., McIntyre, M., Warren, K., Midla, J., Luong, N., and Benedict, K. (2018) Multistate Epidemiology of Histoplasmosis, United States, 2011-2014. Emerg Infect Dis 24, 425-431

4. Antinori, S. (2014) Histoplasma capsulatum: more widespread than previously thought. Am J Trop Med Hyg 90, 982–983

5. Benedict, K., and Mody, R. K. (2016) Epidemiology of Histoplasmosis Outbreaks, United States, 1938-2013. Emerg Infect Dis 22, 370-378

6. Goughenour, K. D., and Rappleye, C. A. (2017) Antifungal therapeutics for dimorphic fungal pathogens. Virulence 8, 211–221

7. Parente-Rocha, J. A., Bailao, A. M., Amaral, A. C., Taborda, C. P., Paccez, J. D., Borges, C. L., and Pereira, M. (2017) Antifungal Resistance, Metabolic Routes as Drug Targets, and New Antifungal Agents: An Overview about Endemic Dimorphic Fungi. Mediators Inflamm 2017, 9870679

8. Pianalto, K. M., and Alspaugh, J. A. (2016) New Horizons in Antifungal Therapy. J Fungi (Basel) 2

9. Gray, K. C., Palacios, D. S., Dailey, I., Endo, M. M., Uno, B. E., Wilcock, B. C., and Burke, M. D. (2012) Amphotericin primarily kills yeast by simply binding ergosterol. Proc Natl Acad Sci U S A 109, 2234–2239

10. Palacios, D. S., Dailey, I., Siebert, D. M., Wilcock, B. C., and Burke, M. D. (2011) Synthesis-enabled functional group deletions reveal key underpinnings of amphotericin B ion channel and antifungal activities. Proc Natl Acad Sci U S A 108, 6733–6738

11. Anderson, T. M., Clay, M. C., Cioffi, A. G., Diaz, K. A., Hisao, G. S., Tuttle, M. D., Nieuwkoop, A. J., Comellas, G., Maryum, N., Wang, S., Uno, B. E., Wildeman, E. L., Gonen, T., Rienstra, C. M., and Burke, M. D. (2014) Amphotericin forms an extramembranous and fungicidal sterol sponge. Nat Chem Biol 10, 400–406

12. Ghannoum, M. A., and Rice, L. B. (1999) Antifungal agents: mode of action, mechanisms of resistance, and correlation of these mechanisms with bacterial resistance. Clin Microbiol Rev 12, 501–517

13. Mor, V., Rella, A., Farnoud, A. M., Singh, A., Munshi, M., Bryan, A., Naseem, S., Konopka, J. B., Ojima, I., Bullesbach, E., Ashbaugh, A., Linke, M. J., Cushion, M., Collins, M., Ananthula, H. K., Sallans, L., Desai, P. B., Wiederhold, N. P., Fothergill, A. W., Kirkpatrick, W. R., Patterson, T., Wong, L. H., Sinha, S., Giaever, G., Nislow, C., Flaherty, P., Pan, X., Cesar, G. V., de Melo Tavares, P., Frases, S., Miranda, K., Rodrigues, M. L., Luberto, C., Nimrichter, L., and Del Poeta, M. (2015) Identification of a New Class of Antifungals Targeting the Synthesis of Fungal Sphingolipids. MBio 6, e00647

14. Rittershaus, P. C., Kechichian, T. B., Allegood, J. C., Merrill, A. H., Jr., Hennig, M., Luberto, C., and Del Poeta, M. (2006) Glucosylceramide synthase is an essential regulator of pathogenicity of Cryptococcus neoformans. J Clin Invest 116, 1651–1659

15. Artunduaga Bonilla, J. J., Honorato, L., Haranahalli, K., Gremiao, I. D. F., Pereira, S. A., Guimaraes, A., Baptista, A. R. S., de, M. T. P., Rodrigues, M. L., Miranda, K., Ojima, I., Del Poeta, M., and Nimrichter, L. (2021) Antifungal activity of Acylhydrazone derivatives against Sporothrix spp. Antimicrob Agents Chemother

16. Dos Reis, T. F., Horta, M. A. C., Colabardini, A. C., Fernandes, C. M., Silva, L. P., Bastos, R. W., Fonseca, M. V. L., Wang, F., Martins, C., Rodrigues, M. L., Silva Pereira, C., Del Poeta, M., Wong, K. H., and Goldman, G. H. (2021) Screening of Chemical Libraries for New Antifungal Drugs against Aspergillus fumigatus Reveals Sphingolipids Are Involved in the Mechanism of Action of Miltefosine. mBio 12, e0145821

17. Cheng, Y. S., Roma, J. S., Shen, M., Mota Fernandes, C., Tsang, P. S., Forbes, H. E., Boshoff, H., Lazzarini, C., Del Poeta, M., Zheng, W., and Williamson, P. R. (2021) Identification of Antifungal Compounds against Multidrug-Resistant Candida auris Utilizing a High-Throughput Drug-Repurposing Screen. Antimicrob Agents Chemother 65

18. Weete, J. D., Abril, M., and Blackwell, M. (2010) Phylogenetic distribution of fungal sterols. PLoS One 5, e10899

19. Vallejo, M. C., Nakayasu, E. S., Longo, L. V., Ganiko, L., Lopes, F. G., Matsuo, A. L., Almeida, I. C., and Puccia, R. (2012) Lipidomic analysis of extracellular vesicles from the pathogenic phase of Paracoccidioides brasiliensis. PLoS One 7, e39463

20. Grandmougin-Ferjani, A., Dalpé, Y., Hartmann, M.-A., Laruelle, F., and Sancholle, M. (1999) Sterol distribution in arbuscular mycorrhizal fungi. Phytochemistry 50, 1027–1031

21. Debieu, D., Corio-Costet, M.-F., Steva, H., Malosse, C., and Leroux, P. (1995) Sterol composition of the vine powdery mildew fungus, Uncinula necator: Comparison of triadimenol-sensitive and resistant strains. Phytochemistry 39, 293–300

22. Weete, J. D., and Gandhi, S. R. (1999) Sterols and fatty acids of the Mortierellaceae: taxonomic implications. Mycologia 91, 642–649

23. Nakayasu, E. S., Nicora, C. D., Sims, A. C., Burnum-Johnson, K. E., Kim, Y. M., Kyle, J. E., Matzke, M. M., Shukla, A. K., Chu, R. K., Schepmoes, A. A., Jacobs, J. M., Baric, R. S., Webb-Robertson, B. J., Smith, R. D., and Metz, T. O. (2016) MPLEx: a Robust and Universal Protocol for Single-Sample Integrative Proteomic, Metabolomic, and Lipidomic Analyses. mSystems 1

24. Beck, M., Schmidt, A., Malmstroem, J., Claassen, M., Ori, A., Szymborska, A., Herzog, F., Rinner, O., Ellenberg, J., and Aebersold, R. (2011) The quantitative proteome of a human cell line. Mol Syst Biol 7, 549

25. Casanovas, A., Sprenger, R. R., Tarasov, K., Ruckerbauer, D. E., Hannibal-Bach, H. K., Zanghellini, J., Jensen, O. N., and Ejsing, C. S. (2015) Quantitative analysis of proteome and lipidome dynamics reveals functional regulation of global lipid metabolism. Chem Biol 22, 412–425

26. Zamith-Miranda, D., Heyman, H. M., Cleare, L. G., Couvillion, S. P., Clair, G. C., Bredeweg, E. L., Gacser, A., Nimrichter, L., Nakayasu, E. S., and Nosanchuk, J. D. (2019) Multi-omics Signature of Candida auris, an Emerging and Multidrug-Resistant Pathogen. mSystems 4

27. Tamaki, H., Shimada, A., Ito, Y., Ohya, M., Takase, J., Miyashita, M., Miyagawa, H., Nozaki, H., Nakayama, R., and Kumagai, H. (2007) LPT1 encodes a membrane-bound O-acyltransferase involved in the acylation of lysophospholipids in the yeast Saccharomyces cerevisiae. J Biol Chem 282, 34288–34298

28. Damude, H. G., Zhang, H., Farrall, L., Ripp, K. G., Tomb, J. F., Hollerbach, D., and Yadav, N. S. (2006) Identification of bifunctional delta12/omega3 fatty acid desaturases for improving the ratio of omega3 to omega6 fatty acids in microbes and plants. Proc Natl Acad Sci U S A 103, 9446–9451

29. Radovanovic, N., Thambugala, D., Duguid, S., Loewen, E., and Cloutier, S. (2014) Functional characterization of flax fatty acid desaturase FAD2 and FAD3 isoforms expressed in yeast reveals a broad diversity in activity. Mol Biotechnol 56, 609–620

30. Deepe, G. S., Jr., and Buesing, W. R. (2012) Deciphering the pathways of death of Histoplasma capsulatum-infected macrophages: implications for the immunopathogenesis of early infection. J Immunol 188, 334–344

31. Metzl-Raz, E., Kafri, M., Yaakov, G., Soifer, I., Gurvich, Y., and Barkai, N. (2017) Principles of cellular resource allocation revealed by condition-dependent proteome profiling. Elife 6

32. Domer, J. E., and Hamilton, J. G. (1971) The readily extracted lipids of Histoplasma capsulatum and Blastomyces dermatitidis. Biochim Biophys Acta 231, 465–478

33. Phetsuksiri, B., Jackson, M., Scherman, H., McNeil, M., Besra, G. S., Baulard, A. R., Slayden, R. A., DeBarber, A. E., Barry, C. E., 3rd, Baird, M. S., Crick, D. C., and Brennan, P. J. (2003) Unique mechanism of action of the thiourea drug isoxyl on Mycobacterium tuberculosis. J Biol Chem 278, 53123-53130

34. Phetsuksiri, B., Baulard, A. R., Cooper, A. M., Minnikin, D. E., Douglas, J. D., Besra, G. S., and Brennan, P. J. (1999) Antimycobacterial activities of isoxyl and new derivatives through the inhibition of mycolic acid synthesis. Antimicrob Agents Chemother 43, 1042–1051

35. Tousek, J. (1970) On the clinical effectiveness of Isoxyl. Antibiot Chemother 16, 149–155

36. Urbancik, B. (1970) Clinical experience with thiocarlide (Isoxyl). Antibiot Chemother 16, 117–123

37. Wang, C., and Hickey, A. J. (2010) Isoxyl particles for pulmonary delivery: In vitro cytotoxicity and potency. Int J Pharm 396, 99–104

38. Maldonado, R. A., Kuniyoshi, R. K., Linss, J. G., and Almeida, I. C. (2006) Trypanosoma cruzi oleate desaturase: molecular characterization and comparative analysis in other trypanosomatids. J Parasitol 92, 1064–1074

39. Alloatti, A., Testero, S. A., and Uttaro, A. D. (2009) Chemical evaluation of fatty acid desaturases as drug targets in Trypanosoma cruzi. Int J Parasitol 39, 985–993

40. Bopp, L. H., Baltch, A. L., Ritz, W. J., Michelsen, P. B., and Smith, R. P. (2006) Antifungal effect of voriconazole on intracellular Candida glabrata, Candida krusei and Candida parapsilosis in human monocyte-derived macrophages. J Med Microbiol 55, 865–870

41. Chattopadhyay, P., Banerjee, S. K., Sen, K., and Chakrabarti, P. (1985) Lipid profiles of Aspergillus niger and its unsaturated fatty acid auxotroph, UFA2. Can J Microbiol 31, 352-355

42. Chattopadhyay, P., Banerjee, S. K., Sen, K., and Chakrabarti, P. (1987) Lipid profiles of conidia of Aspergillus niger and a fatty acid auxotroph. Can J Microbiol 33, 1116–1120

43. Pettersen, R. J., Salem, M., Skorve, J., Ulvik, R. J., Berge, R. K., and Nordrehaug, J. E. (2008) Pharmacology and safety of tetradecylthioacetic acid (TTA): phase-1 study. J Cardiovasc Pharmacol 51, 410–417

44. Kluepfel, D., Bagli, J., Baker, H., Charest, M. P., and Kudelski, A. (1972) Myriocin, a new antifungal antibiotic from Myriococcum albomyces. J Antibiot (Tokyo) 25, 109–115

45. Perdoni, F., Signorelli, P., Cirasola, D., Caretti, A., Galimberti, V., Biggiogera, M., Gasco, P., Musicanti, C., Morace, G., and Borghi, E. (2015) Antifungal activity of Myriocin on clinically relevant Aspergillus fumigatus strains producing biofilm. BMC Microbiol 15, 248

46. de Melo, N. R., Abdrahman, A., Greig, C., Mukherjee, K., Thornton, C., Ratcliffe, N. A., Vilcinskas, A., and Butt, T. M. (2013) Myriocin significantly increases the mortality of a non-mammalian model host during Candida pathogenesis. PLoS One 8, e78905

47. Strader, C. R., Pearce, C. J., and Oberlies, N. H. (2011) Fingolimod (FTY720): a recently approved multiple sclerosis drug based on a fungal secondary metabolite. J Nat Prod 74, 900–907

48. Caretti, A., Torelli, R., Perdoni, F., Falleni, M., Tosi, D., Zulueta, A., Casas, J., Sanguinetti, M., Ghidoni, R., Borghi, E., and Signorelli, P. (2016) Inhibition of ceramide de novo synthesis by myriocin produces the double effect of reducing pathological inflammation and exerting antifungal activity against A. fumigatus airways infection. Biochim Biophys Acta 1860, 1089–1097

49. Lazzarini, C., Haranahalli, K., Rieger, R., Ananthula, H. K., Desai, P. B., Ashbaugh, A., Linke, M. J., Cushion, M. T., Ruzsicska, B., Haley, J., Ojima, I., and Del Poeta, M. (2018) Acylhydrazones as Antifungal Agents Targeting the Synthesis of Fungal Sphingolipids. Antimicrob Agents Chemother 62

50. Gao, J., Wang, H., Li, Z., Wong, A. H., Wang, Y. Z., Guo, Y., Lin, X., Zeng, G., Liu, H., Wang, Y., and Wang, J. (2018) Candida albicans gains azole resistance by altering sphingolipid composition. Nat Commun 9, 4495

51. Spitzer, M., Griffiths, E., Blakely, K. M., Wildenhain, J., Ejim, L., Rossi, L., De Pascale, G., Curak, J., Brown, E., Tyers, M., and Wright, G. D. (2011) Cross-species discovery of syncretic drug combinations that potentiate the antifungal fluconazole. Mol Syst Biol 7, 499

52. Cleare, L. G., Zamith, D., Heyman, H. M., Couvillion, S. P., Nimrichter, L., Rodrigues, M. L., Nakayasu, E. S., and Nosanchuk, J. D. (2020) Media matters! Alterations in the loading and release of Histoplasma capsulatum extracellular vesicles in response to different nutritional milieus. Cell Microbiol, e13217

53. Gietz, R. D., Schiestl, R. H., Willems, A. R., and Woods, R. A. (1995) Studies on the transformation of intact yeast cells by the LiAc/SS-DNA/PEG procedure. Yeast 11, 355–360

54. Matos Baltazar, L., Nakayasu, E. S., Sobreira, T. J., Choi, H., Casadevall, A., Nimrichter, L., and Nosanchuk, J. D. (2016) Antibody Binding Alters the Characteristics and Contents of Extracellular Vesicles Released by Histoplasma capsulatum. mSphere 1

55. Albuquerque, P. C., Nakayasu, E. S., Rodrigues, M. L., Frases, S., Casadevall, A., Zancope-Oliveira, R. M., Almeida, I. C., and Nosanchuk, J. D. (2008) Vesicular transport in Histoplasma capsulatum: an effective mechanism for trans-cell wall transfer of proteins and lipids in ascomycetes. Cell Microbiol 10, 1695–1710

56. Burnet, M. C., Zamith-Miranda, D., Heyman, H. M., Weitz, K. K., Bredeweg, E. L., Nosanchuk, J. D., and Nakayasu, E. S. (2020) Remodeling of the Histoplasma Capsulatum Membrane Induced by Monoclonal Antibodies. Vaccines (Basel) 8

57. Wilhelm, M., Schlegl, J., Hahne, H., Gholami, A. M., Lieberenz, M., Savitski, M. M., Ziegler, E., Butzmann, L., Gessulat, S., Marx, H., Mathieson, T., Lemeer, S., Schnatbaum, K., Reimer, U., Wenschuh, H., Mollenhauer, M., Slotta-Huspenina, J., Boese, J. H., Bantscheff, M., Gerstmair, A., Faerber, F., and Kuster, B. (2014) Mass-spectrometry-based draft of the human proteome. Nature 509, 582–587

58. Baltazar, L. M., Zamith-Miranda, D., Burnet, M. C., Choi, H., Nimrichter, L., Nakayasu, E. S., and Nosanchuk, J. D. (2018) Concentration-dependent protein loading of extracellular vesicles released by Histoplasma capsulatum after antibody treatment and its modulatory action upon macrophages. Sci Rep 8, 8065

59. Dautel, S. E., Kyle, J. E., Clair, G., Sontag, R. L., Weitz, K. K., Shukla, A. K., Nguyen, S. N., Kim, Y. M., Zink, E. M., Luders, T., Frevert, C. W., Gharib, S. A., Laskin, J., Carson, J. P., Metz, T. O., Corley, R. A., and Ansong, C. (2017) Lipidomics reveals dramatic lipid compositional changes in the maturing postnatal lung. Sci Rep 7, 40555

60. Kyle, J. E., Crowell, K. L., Casey, C. P., Fujimoto, G. M., Kim, S., Dautel, S. E., Smith, R. D., Payne, S. H., and Metz, T. O. (2017) LIQUID: an-open source software for identifying lipids in LC-MS/MS-based lipidomics data. Bioinformatics 33, 1744–1746

61. Pluskal, T., Castillo, S., Villar-Briones, A., and Oresic, M. (2010) MZmine 2: modular framework for processing, visualizing, and analyzing mass spectrometry-based molecular profile data. BMC Bioinformatics 11, 395

62. Kim, Y. M., Schmidt, B. J., Kidwai, A. S., Jones, M. B., Deatherage Kaiser, B. L., Brewer, H. M., Mitchell, H. D., Palsson, B. O., McDermott, J. E., Heffron, F., Smith, R. D., Peterson, S. N., Ansong, C., Hyduke, D. R., Metz, T. O., and Adkins, J. N. (2013) Salmonella modulates metabolism during growth under conditions that induce expression of virulence genes. Mol Biosyst 9, 1522–1534

63. Hiller, K., Hangebrauk, J., Jager, C., Spura, J., Schreiber, K., and Schomburg, D. (2009) MetaboliteDetector: comprehensive analysis tool for targeted and nontargeted GC/MS based metabolome analysis. Anal Chem 81, 3429–3439

64. Kind, T., Wohlgemuth, G., Lee, D. Y., Lu, Y., Palazoglu, M., Shahbaz, S., and Fiehn, O. (2009) FiehnLib: mass spectral and retention index libraries for metabolomics based on quadrupole and time-of-flight gas chromatography/mass spectrometry. Anal Chem 81, 10038–10048

65. Hartmann, A., and Jozefowicz, A. M. (2018) VANTED: A Tool for Integrative Visualization and Analysis of -Omics Data. Methods Mol Biol 1696, 261–278

66. Munshi, M. A., Gardin, J. M., Singh, A., Luberto, C., Rieger, R., Bouklas, T., Fries, B. C., and Del Poeta, M. (2018 The Role of Ceramide Synthases in the Pathogenicity of *Cryptococcus neoformans*. Cell reports 22, 1392–1400

